# Mild drought induces phenotypic and DNA methylation plasticity but no transgenerational effects in Arabidopsis

**DOI:** 10.1101/370320

**Authors:** Tom JM Van Dooren, Amanda Bortolini Silveira, Elodie Gilbault, José M. Jiménez-Gómez, Antoine Martin, Liên Bach, Sébastien Tisné, Leandro Quadrana, Olivier Loudet, Vincent Colot

## Abstract

Whether environmentally induced changes in phenotypes can be heritable is a topic with revived interest, in part because of observations in plants that heritable trait variation can occur without DNA sequence mutations. This other system of inheritance, called transgenerational epigenetics, typically involves differences in DNA methylation that are stable across multiple generations. However, it remains unknown if such a system responds to environmental changes and if it could therefore provide a rapid way for plants to generate adaptive heritable phenotypic variation. Here, we used a well-controlled phenotyping platform and whole-genome bisulfite sequencing to investigate potential heritable effects of mild drought applied over two successive generations in *Arabidopsis thaliana*. Plastic phenotypic responses were observed in plants exposed to drought. After an intervening generation without stress, descendants of stressed and non-stressed plants were phenotypically indistinguishable, except for very few trait-based parental effects, and irrespective of whether they were grown in control conditions or under water deficit. Moreover, while mild drought induced changes to the DNA methylome of exposed plants, DNA methylation variants were not inherited. These findings add to the growing body of evidence indicating that transgenerational epigenetics is not a common response of plants to environmental changes.

## Introduction

It is now well established that DNA mutations are not the only source of heritable phenotypic variation in plants. An additional system of inheritance, often referred to as transgenerational epigenetics, typically involves stable differences of DNA methylation at or near transposable element (TE) sequences that are adjacent to genes (Quadrana & Colot, 2016). In the reference plant Arabidopsis, most TE sequences are methylated at all cytosines, with methylation levels generally highest at CG sites (>80%), intermediate at CHG sites (40-60%) and lowest at CHH sites (<20%) (Cokus *et al.*, 2008; Lister *et al.*, 2008). TE sequences are methylated as a result of the combined activity of multiple DNA methyltransferases (Law & Jacobsen, 2010; Stroud *et al.*, 2013; Stroud *et al.*, 2014) and can also be actively demethylated by DNA glycosylases, which excise methylated cytosines from DNA (Law & Jacobsen, 2010). Demethylation is most pronounced in the central cell and leads after fertilization to a global hypomethylation of TE sequences in the endosperm, specifically on the maternally-derived chromosomes (Satyaki & Gehring, 2017). In contrast, because methylation dynamics is restricted to CHG and especially CHH sites (Bouyer *et al.*, 2017; Kawakatsu *et al.*, 2017; Lin *et al.*, 2017), TE sequences remain highly methylated in the female and male germlines as well as the embryo. In turn, this limited reprogramming of DNA methylation patterns between generations implies a considerable potential genome-wide for transgenerational epiallelic variation following accidental loss of DNA methylation. However, because the *de novo* DNA methylation machinery targets distinct TE sequences with varying efficiency (Teixeira *et al.*, 2009; Zemach *et al.*, 2013), this potential is in fact not uniformly distributed among TE-containing alleles. Thus, while experimentally induced epiallelic variation can persist for at least 8 generations and presumably many more at some TE-containing loci, it is fully erased within one or a few generations at others (Johannes *et al.*, 2009; Teixeira *et al.*, 2009; Colome-Tatche *et al.*, 2012).

Being sessile organisms, plants are often exposed to environmental stresses, which can have a profound impact on the growth and development of not only the exposed individuals but also their offspring. These parental (typically maternal) effects are well documented (Blödner *et al.*, 2007; Galloway & Etterson, 2007; Donohue, 2009; Herman & Sultan, 2011; Crisp *et al.*, 2016; Van Dooren *et al.*, 2016). In contrast, few experimental studies have been conducted to determine if a phenotypic memory of environmental stress can persist over multiple successive generations. In one early study, genetically identical Arabidopsis lines grown under mild heat during the reproductive phase (from bolting onward) over two generations and then grown under normal conditions for one more generation produced progeny with an ameliorated response to heat treatment compared to control progeny derived from non-treated lines (Whittle *et al.*, 2009). However, as heat was applied during reproductive growth, it affected the gametes and the developing seeds produced by the treated plants (Whittle *et al.*, 2009). Therefore, parental effects could still be responsible for the ameliorated response to heat seen in the progeny of these individuals (Blödner *et al.*, 2007; Pecinka & Mittelsten Scheid, 2012). Consistent with this possibility, another study indicated that when heat stress was applied during vegetative growth only, phenotypic effects did not persist for more than one generation (Suter & Widmer, 2013b; Suter & Widmer, 2013a). Similarly, when Arabidopsis plants were infected with pathogens, increased resistance was reported in the immediate progeny as well as in the second generation, but only when infections were carried out until the reproductive phase (Boyko *et al.*, 2010; Luna *et al.*, 2012; Slaughter *et al.*, 2012). Salt stress memory across generations was also investigated and findings all point to an absence of *bona fide* transgenerational effects (Boyko *et al.*, 2010; Suter & Widmer, 2013b; Suter & Widmer, 2013a;Groot *et al.*, 2016; Wibowo *et al.*, 2016). Thus, the moment when stress is experienced in the plant life history matters for the strength of the effect, its transmission across generations and probably the causation of the effects. Finally, in the few cases where this was looked at, DNA methylation changes were observed in response to stress and some of these changes were transmitted, but again transmission was limited to the immediate progeny only (Secco *et al.*, 2015; Wibowo *et al.*, 2016; Ganguly *et al.*, 2017).

Here, we set out to determine if prolonged water deficit, a common stress that plants face in natural settings, could lead to new or altered transgenerational effects. Our experimental design and analysis combine several distinctive key aspects compared to previous studies on responses to stress. First, we tested four accessions (Shahdara, Bur-0, Tsu-0 and Cvi-0) in addition to the reference accession Col-0, in order to assess how stress response, possibly in relation to distinct TE landscapes, could influence phenotypic patterns. Second, we used a well-controlled multigenerational experimental design where the magnitude and type of environmental stress was replicated across generations for all five accessions. This design enabled us to readily distinguish parental from transgenerational effects and to investigate interactions of maternal effects and plasticity. Third, we used the robotic platform Phenoscope, which ensures uniform conditions during vegetative growth and enables continuous as well as precise phenotype tracking. Fourth, we analyzed the development of phenotypes over time, and estimate magnitudes of trait-based maternal effects for five traits in combination with plasticity. Fifth, we carried out an in-depth assessment of the likelihood of transgenerational epigenetics at the DNA methylation level in the reference accession Col-0. Our results indicate that mild drought induces phenotypic plasticity in each of the five accessions, but does not lead to any significant change in terms of heritable effects. In addition, we obtained methylome data at single cytosine resolution for Col-0, which again indicate that mild drought induces intragenerational DNA methylation changes only, which are restricted to CHH sites and affect TE sequences predominantly. Taken together, our findings add to the growing body of evidence indicating that plants do not commonly generate transgenerational effects in response to changes in the environment.

## Material and methods

### Plant material and growth conditions

To investigate interactions between genotype and environment (G×E) in the response to mild drought (Bouchabke *et al.*, 2008), we considered the five accessions Col-0 (Col; Versailles stock # 186AV), Shahdara (Sha; 236AV), Bur-0 (Bur; 172AV), Tsu-0 (Tsu; 91AV) and Cvi-0 (Cvi; 166AV), which were obtained from the Versailles stock center (http://publiclines.versailles.inra.fr/). The four non-reference accessions were chosen because they show similar flowering time to Col-0 but extensive nucleotide as well as DNA methylation divergence among themselves and with Col-0 (Consortium, 2016; Kawakatsu *et al.*, 2016a). Isogenic lines for each accession were grown under well-watered control and mild drought stress conditions for three generations (Fig. 1), using the robotic platform Phenoscope (https://phenoscope.versailles.inra.fr/), which ensures uniform conditions during vegetative growth and enables precise phenotype tracking (Tisne *et al.*, 2013). At the first generation (G1), 12 individuals (descending from the same mother plant) per accession and per treatment were grown and half were used to establish 6 independent founder lines, which were then maintained throughout the experiment by single seed descent propagation. In the following generations, six replicates per accession, treatment and trajectory (life history) were grown, with the exception of the third generation (G3) where only four replicates were grown due to space limitations on the phenotyping robot (Fig. 1). Growth conditions (control and mild drought stress) in each generation were as described in detail elsewhere (Tisne *et al.*, 2013). Briefly, seeds were stratified for 4 days in the dark at 4°C and germinated for 8 days on peat moss plugs before the plugs were transferred to the robot. Individual plants were then cultivated on the Phenoscope for another 21 days under short days in order to minimize developmental differences between accessions and delay flowering transition. The first week, during germination, soil was saturated with water. During the first 7 days of growth on the Phenoscope (Day 9 to 15 after sowing), soil water content gradually decreased through controlled watering until it reached either 60% (control) or 30% (stress), which was then strictly maintained at this level for the remaining 2 weeks (until Day 29 after sowing). At the end of the Phenoscope experiment, plants were moved to a standard growth chamber with optimal watering and long-day conditions to allow for flowering and seed production. This strategy ensured that gametes or seeds were not themselves exposed to mild drought.

**Fig. 1.**
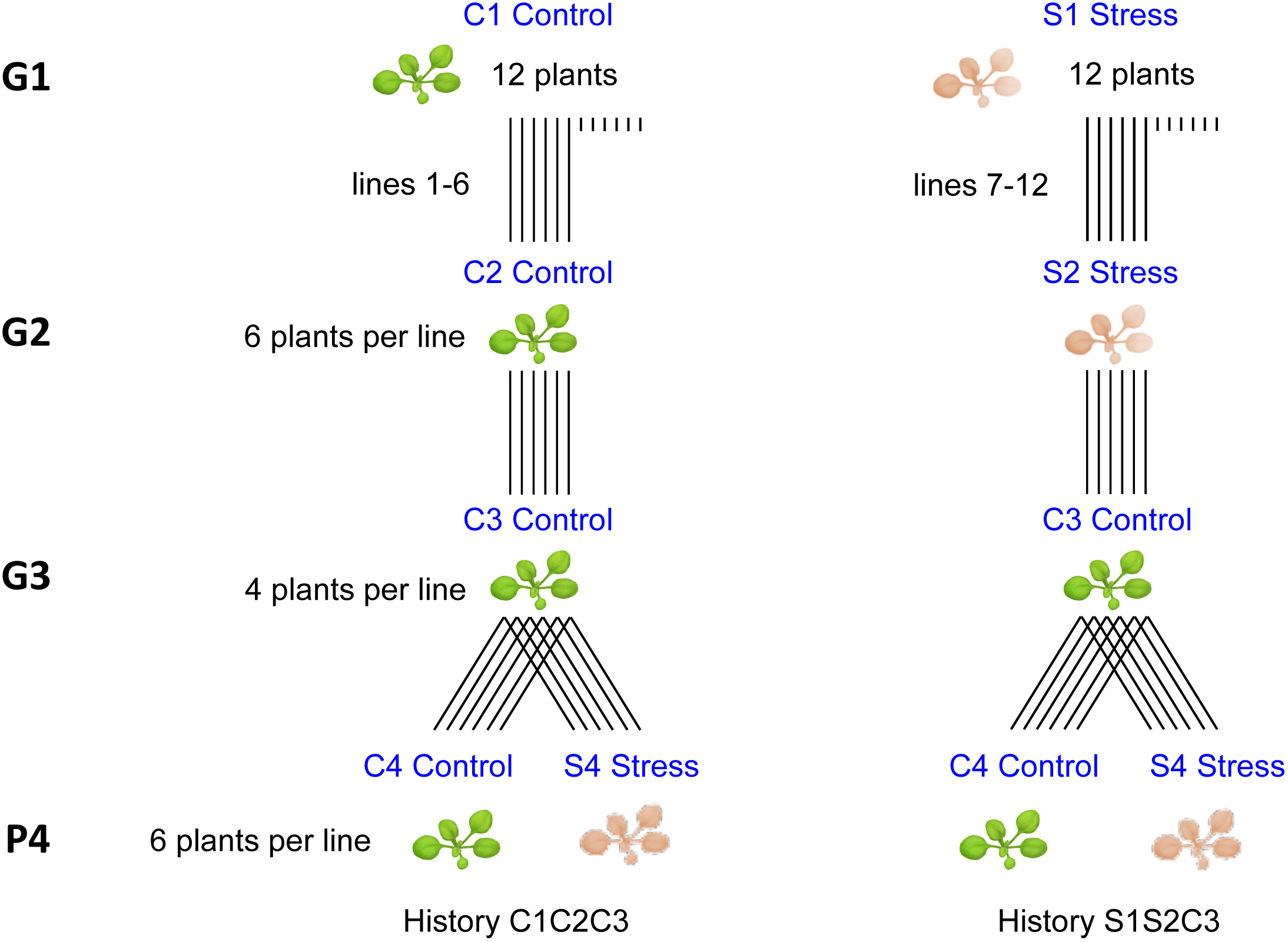
Schematic representation of the multigenerational experimental design. G, generation of growth; P, phenotyping experiment.

For the subsequent generation, collected seeds were sieved so as to avoid sowing seeds that were in the top 10% size or the bottom 10% of each line’s seed size distribution. Seed size range varied between accessions (with Cvi and Bur having bigger seeds than Col and Sha especially), but not between lines within accessions. A final phenotyping experiment (P4; Fig. 1) was conducted on lines from two selected trajectories (S1S2C3 versus C1C2C3).

### Phenotyping

At every generation, zenithal rosette images of each individual plant were taken daily and segmented as described previously (Tisne *et al.*, 2013) to extract projected rosette area (PRA; a good proxy for rosette biomass at these development stages), rosette radius (the radius of the circle encompassing the rosette), compactness (the ratio between PRA and the rosette circle area) as well as the red, green and blue components of the segmented rosette image. We report our phenotypic analysis of the last generation of plants grown on the Phenoscope (P4).

### Size and relative growth analysis

Plant cohorts might contain different groups with different properties and responses to treatments, while group membership might be unknown for each individual. To investigate such large initial heterogeneity among the plants selected for growth on the robot, initial PRA distributions on Day 9 after sowing (essentially the summed cotyledon areas) were inspected by finite mixture analysis using the FlexMix library (Grun & Leisch, 2007). Gaussian models for initial log(PRA) with different numbers of component distributions (1 to 3) and different fixed treatment effects (stress/control in P4, stress/control in G1 and G2, i.e. G1/G2) for each component were compared using three information criteria (AIC, BIC and ICL). The model with the lowest values of the information criteria was chosen each time, with preference given to the ranking of ICL in case of discordance. Significant fixed effects are reported (*z*-test) for preferred models, where these occur.

Initial and final values of log(PRA) were further studied in detail using linear mixed models (Pinheiro & Bates, 2000). The maximal models fitted contained random line effects with different variances in control and stress groups in G1/G2. Models with more involved random effect specifications did not converge. Fixed effects were the exposure to stress in G1/G2, stress treatment in P4, pot order effects on the robot and interactions of these variables. The maximal model contained heterogeneous error variances, different for each control/stress combination in G1/G2 and P4. Model comparisons and simplifications were carried out using likelihood ratio tests LRT (using a REML fit for random effects, ML for fixed). Non-significant effects were removed one by one. Simplifications were first attempted in the random effects, then in the error variances and after that in the fixed effects, starting with the highest order interactions. Tests and estimates for random effects and error variances are reported for a REML model containing all fixed effects. P-values for fixed effects are reported with respect to the minimum adequate model (MAM) selected, i.e., the first model encountered that only contains significant effects.

We also analyzed relative growth rates of PRA to understand the gradual response to environmental conditions. We first inspected the per-day relative increase of log(PRA) using generalized additive mixed models (gamms; (Wood, 2017) and subsequently fitted mixed linear models (Pinheiro & Bates, 2000) to growth rates in age intervals that appeared linear. Per separate treatment combination and accession, a gamm was fitted with smooths for day (age) and pot order effects (pot number). The gamm’s assume random variation between individuals and exponentially decaying correlations between observations on the same individual. The mixed linear models fitted to restricted age intervals contained the same variables as the models for final size above, with the addition of random variation between individuals within lines, fixed age (day) effects and interactions of age with the other fixed effects.

### Phenotypic maternal trait-based effects

Our analysis of PRA is detailed and accounts for effects of ancestral environments in G1/G2, a potentially transmitted maternal environmental effect, but does not include trait-based maternal effects (Kirkpatrick & Lande, 1989). To investigate these effects and whether ancestral environments (i.e. memory) affect their strength and transmission, we analyzed all traits recovered from digital images in a manner that could be applied to all traits equally. We restricted the analysis to the set of traits measured in both G3 and P4 and with correlations between them in P4 below 0.9 (for instance the image green component mean was too correlated to the red component mean to be included as well). Per accession, we thus fitted linear mixed effects models to the log-transformed trait values after day 23. This part of the age trajectory of these traits was always approximately linear. For all traits, we tested whether trait-based parental effects were present and differed between treatments in G1/G2 (ancestral environment) and P4 (plasticity). We used maternal trait values on day 29 of G3 as explanatory variables to model the trait-based effects. We removed data on a few plants with outlying patterns for the increase in log(PRA) before analysis: Observations with a Cook’s distance value (in a simple regression on age) that was larger than one over the number of observations were removed. We fitted maximal mixed models to the data per accession and per trait with random effects of line and individual and heterogeneous error variances that could all differ between environmental treatments experienced for that line in G1/G2 and P4. The model contained fixed effects of the G1/G2 and P4 environmental treatments, pot order effects, age effects, the effects of the maternal trait values for that line in G3 (difference from the overall mean) and interactions of these (except for age×trait interactions and interactions between traits). Note that we thus test whether strengths of maternal trait-based effects depend on environmental conditions. Model selection was carried out as above. However, we observed that selected models often had confidence intervals for the maternal effect slopes that still overlapped with zero or with each other and we simplified such effects out of the models. To interpret the results more easily and to have a graphical means to assess the validity of mixed model predictions, we also fitted linear regressions to offspring trait – maternal trait combinations. All statistical analyses were conducted using R (Team, 2005).

### Whole-genome bisulfite sequencing (WGBS)

To investigate the impact of mild drought on genomic DNA methylation patterns, WGBS was performed on pooled DNA extracted at day 29 after sowing from mature leaves of 12 Col plants that were being subjected to control or water deficit treatments and on 10 day-old seedlings derived from 5 independent C_1_C_2_ and S_1_S_2_ G2 lines grown under standard *in vitro* conditions. MethylC-seq library preparation and sequencing was performed by BGI (Shenzhen, China) using standard Illumina protocols. Adapter and low-quality sequences were trimmed using Trimming Galore v0.3.3. Mapping was performed on TAIR10 genome annotation using Bismark v0.14.2 (Krueger & Andrews, 2011) and the parameters: --bowtie2, -N 1, -p 3 (alignment); --ignore 5 --ignore_r2 5 --ignore_3prime_r2 1 (methylation extractor). Only uniquely mapping reads were retained. The methylKit package v0.9.4 (Akalin *et al.*, 2012) was used to calculate differential methylation in individual positions (DMPs) or in 100 bp non-overlapping windows (DMRs). Significance of calculated differences was determined using Fisher’s exact test and Benjamin-Hochberg (BH) adjustment of *p*-values (FDR < 0.05) and methylation difference cutoffs of 40% for CG, 20% for CHG and 20% for CHH. Differentially methylated windows within 100bp of each other were merged to form larger DMRs. Cytosine positions covered by more than 100 reads were not considered. For DMP analysis only cytosines covered by a minimum of 6 (CG and CHG) and 10 (CHH) reads in all libraries were considered. Bisulfite conversion rates were estimated by the number of methylated cytosine calls in the chloroplast genome.

### RNA-seq

To investigate the impact of mild drought on gene expression, we performed RNA-seq on leaves isolated from stressed and non-stressed Col-0 plants. Leaf tissue was collected at 23 days after sowing from three Col-0 individuals grown on the Phenoscope under control and mild drought conditions and from the same seed batch as the founding Col-0 individuals used in the transgenerational design. Total RNA was extracted using the Qiagen RNAeasy extraction kit and sequenced at the Genome Center of the Max Planck Institute for Plant Breeding Research in Cologne, Germany. RNA-seq libraries were constructed using the standard Illumina Truseq protocol and sequenced in an Illumina Hiseq 2500 machine. Between 18.3 and 23.7 million reads were obtained per sample (average of 20.7) and aligned to the TAIR10 reference genome using TopHat2 with default parameters (Kim *et al.*, 2013). Reads aligning to multiple locations were removed using samtools’ view with parameter -q 5 (Li *et al.*, 2009). After this filter, between 95.5 and 96.8 percent of the obtained reads were aligned to the reference genome. The number of reads per transcript was counted using the Bioconductor packages Rsamtools and ShortRead (Morgan *et al.*, 2009). Differential expression between samples in control and drought conditions was calculated with the DEseq2 package in R (Love *et al.*, 2014). Genes with q-values lower than 0.05 and log2FC above 0.5 were considered as differentially expressed. TE differential expression was analyzed using TETOOLs (Lerat *et al.*, 2016).

RNA-seq and MethylC-seq sequencing data have been deposited in the ENA short read archive under project number PRJEB27682.

## Results

### Mild drought induces immediate phenotypic plasticity

Growth dynamics of the projected rosette area (PRA) in each generation where we imposed mild drought indicated clear phenotypic plasticity in response to mild drought for the five accessions analyzed, as expected (Tisne *et al.*, 2013). Indeed PRA decreased significantly when accessions were grown under mild drought (generation G1, Fig. 2a) and reached values that are on average 27% to 40% lower than in control conditions at an age of 29 days after sowing, depending on the accession (G1, Fig. 2b).

**Fig. 2.**
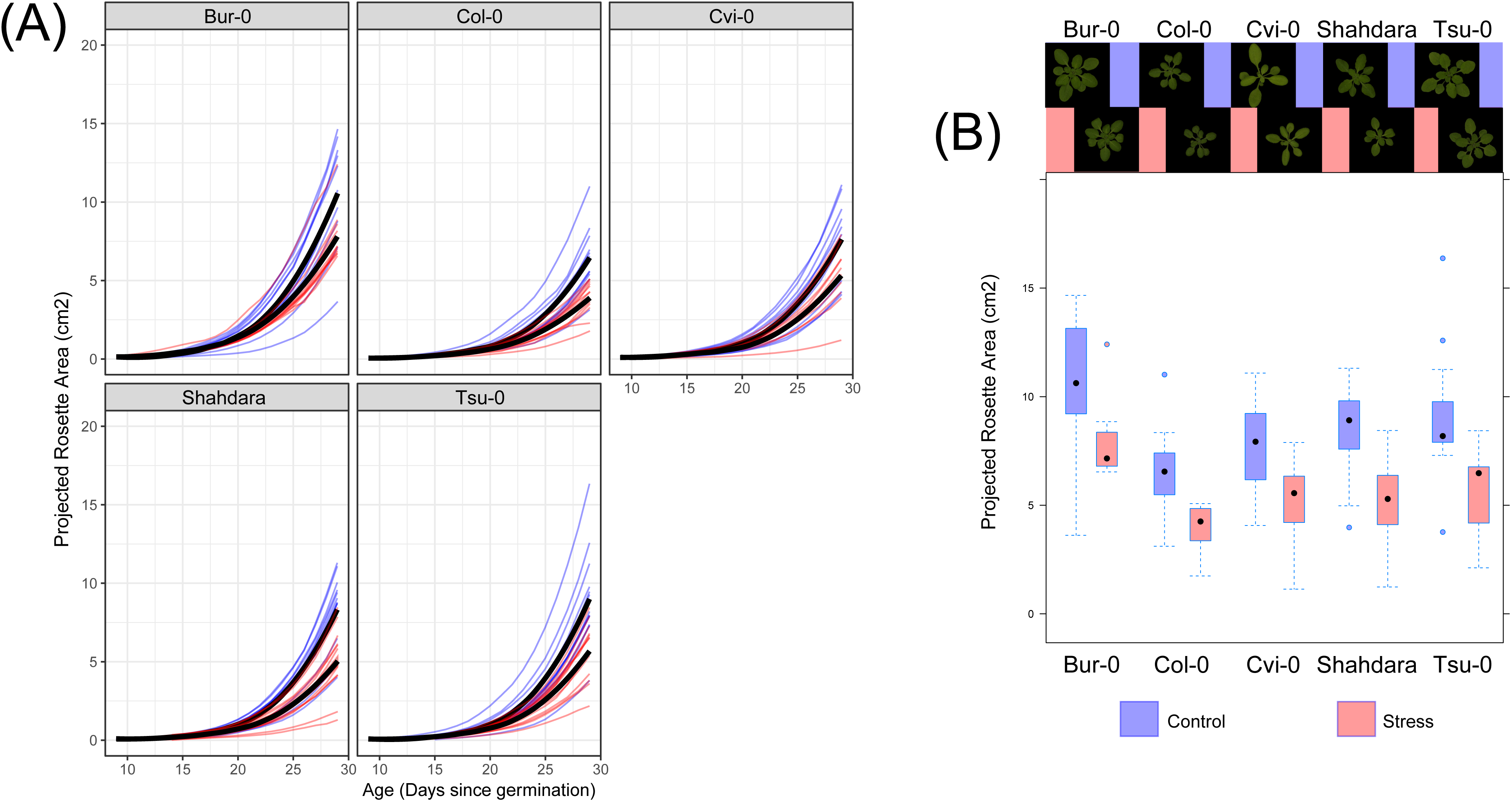
Descriptive analysis of growth curves for projected rosette area (PRA) in the first generation (G1) for the five accessions grown under control or drought (stress) conditions. Individuals that remained below 1 cm2 PRA by the end of the experiment or that died prematurely are not shown. (a) Kinetics of shoot size estimated by daily measurements of PRA. Growth curves of plants in control conditions are in blue, those that experienced mild drought (stress) in red. Black lines represent the averages per group. (b) Box and whiskers plot showing PRA on day 29 for the plants in each accession × treatment combination. Inset photographs show representative plants per accession × treatment.

We then compared, in as much detail as possible, phenotypic traits between the progeny of plants whose parents experienced stress treatments for two consecutive generations and the progeny of plants whose parents never experienced stress (Fig. 1; phenotyping at P4 compares C_1_C_2_C_3_ vs. S_1_S_2_C_3_ trajectories). In both cases one final generation without a stress treatment was included (C3), in order to detect only effects with some capacity to persist independently of the presence of the environmental cue, and to remove direct effects of maternal environments.

We did not find any effect of exposure to stress in the first two generations on initial and final log(PRA) of the progeny of the third generation (P4). The finite mixture analysis of initial log(PRA) indicated that the cohort is not composed of very different groups responding differently to treatments. There are no hidden large heterogeneities among the plants installed on the robot and for each accession, a single Gaussian component was always preferred. In the case of Col, the preferred model did contain an effect of exposure to stress in G1/G2 on average initial size in P4 (*p* = 0.004), with descendants of stressed plants having larger sizes (log(PRA) difference 0.063, s.e. 0.022). We found no effects of exposure to stress in G1/G2 nor P4 on mean initial log(PRA) when taking line variation and pot order effects into account in any of the accessions, hence the single significant effect in the mixture analysis should be attributed to ignoring non-independence in the data. Unexpectedly, we found effects of stress in P4 on initial log(PRA) heterogeneity (residual variance) of two accessions in that generation (Supporting Information Table S1: Cvi and Bur). These effects must be spurious (sampling effect), as the stress treatment has not started yet at this stage. Therefore, we do model variance heterogeneity throughout but do not present nor interpret the results. For mean final size, we find that all accessions except Sha show significant plasticity - their sizes are larger in the control (Table 1).

**Table 1.** Effects of environmental states in G1/G2 and P4 on the mean final projected rosette area log(PRA) in generation P4. Confidence interval results from linear mixed models per accession, based on parameter estimates of minimum adequate models are shown. Symbol "*" indicates accessions where the variance between lines is retained in the model. LRT = Likelihood Ratio Test.

| Accession | Memory of stress in G1/G2 | LRT | Plasticity P4 (Control - Stress) | LRT |
| --- | --- | --- | --- | --- |
| COL | - | NS | [0.197 - 0.375] | p < 0.001 |
| BUR | - | NS | [0.317 - 0.546] | p < 0.001 |
| CVI* | - | NS | [0.394 - 0.557] | p < 0.001 |
| TSU | - | NS | [0.250 - 0.384] | p < 0.001 |
| SHA | - | NS |  | NS |

Our gamm analysis indicated that relative growth rates (RGR) from days 13 to 16 and from days 25 to 28 could be considered linear per individual (Figure 3, representative results for accession Col). Notably, growth rate plasticity in response to drought is not permanent. Indeed, relative growth rates near day 28 are very similar under control and stress conditions, though obviously absolute plant size is smaller.

**Fig. 3.**
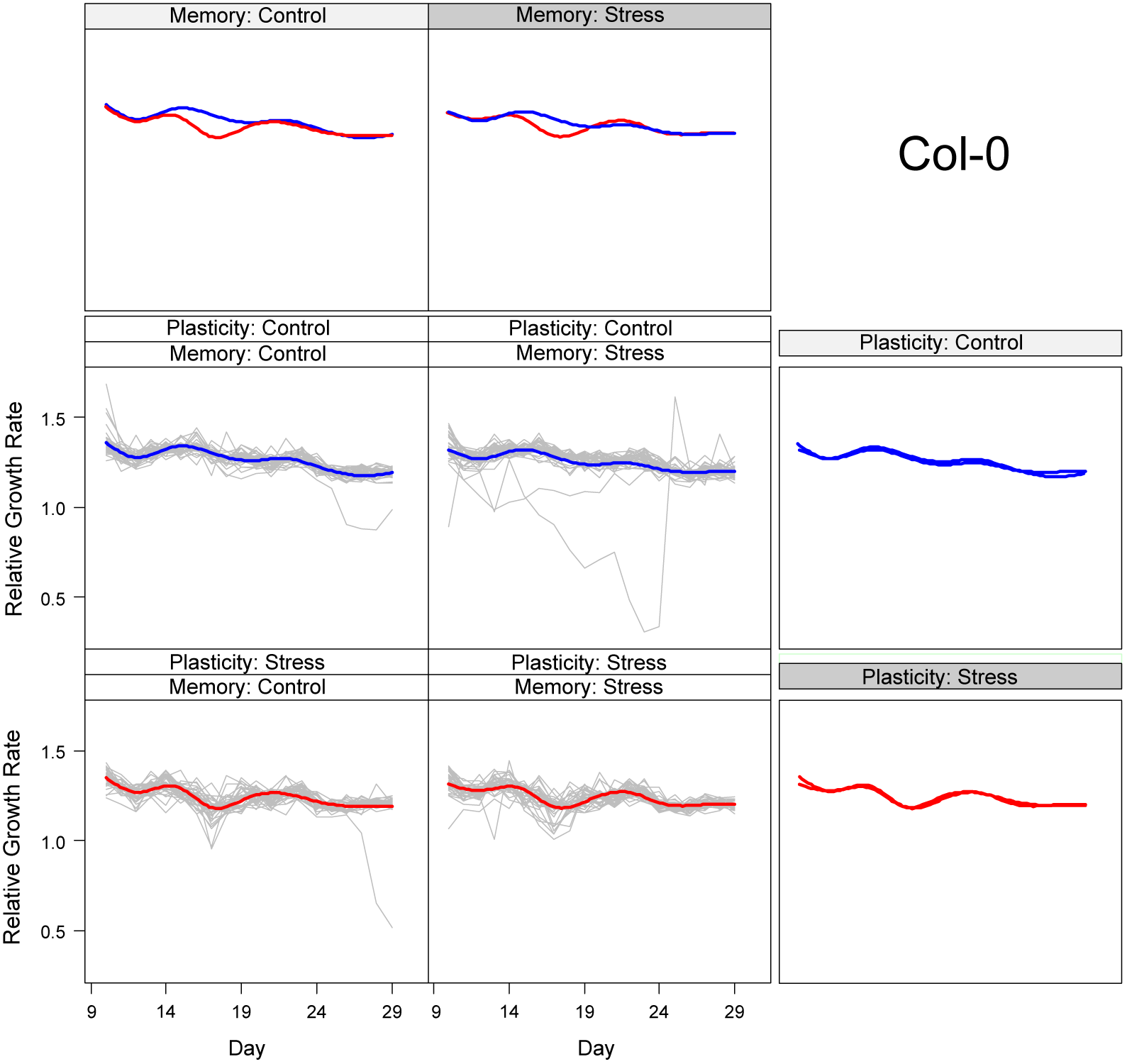
The time-dependent pattern of relative growth rates as predicted for the first pot on the robot (the model accounts for pot order) and for accession Col-0 based on generalized additive mixed models (gamms). On the x-axis the age of the plants in days is given. Plants are nine days old after sowing when the recording starts. Four panels show the raw data per treatment combination. Above and to the right of each column and row, predicted trajectories are grouped for two panels each time so that pairwise comparisons between G1/G2 (Memory) and P4 (Plasticity) treatment levels can be made. The raw data values are shown in grey together with a predicted relative growth rate trajectory per combination of treatments in G1/G2 (Memory) and P4 (Plasticity). P4 treatments are shown in blue (Control) or black (Stress). The graphs demonstrate clear growth plasticity in response to mild drought and that plants manage to compensate the initial drop in relative growth rate shortly after the mild drought stress has reached a stable level at day 20. Note that there are very few plants with outlying patterns and that these have very low growth rates for a restricted age window only.

Modeling RGR demonstrates a steeper decrease from five to eight days after the water supply is reduced in all accessions (Table 2, Fig. 3) and a corresponding decrease in the growth rate intercept for the control group (a more steeply decreasing function has a more positive intercept at day zero as a side effect). However, this initial decrease is followed by a recovery. For three out of five accessions, plants subjected to mild drought acclimate and recover RGR similar to that of control plants (days 25 to 28, Table 3, Fig. 3). For Cvi the recovery is incomplete and for Sha we find some compensatory growth; RGR decreases less with age in the stress group. There are no memory effects of exposure to stress in G1/G2 on average growth rate values in P4. Different magnitudes of between-individual variance across the trajectories do occur for Col, Cvi and Bur. The individual variation in relative growth rates is larger from days 13 to 16 for the Col individuals that descend from parent that experienced stress in G1/G2 (*p* < 0.001); for Bur the variance in the descendants of G1/G2 stress group is smaller (*p* = 0.022). For Col, the growth rate variance from days 25 to 28 is smaller for the descendants of the G1/G2 stress group (*p* < 0.001). For Cvi, this variance is larger in the descendants of the G1/G2 stress group (*p* < 0.001). This pattern points out that amounts of trait variation at the end of an experiment or at the moment where selection occurs could be determined by intricate time-dependent variances in growth processes.

**Table 2.**
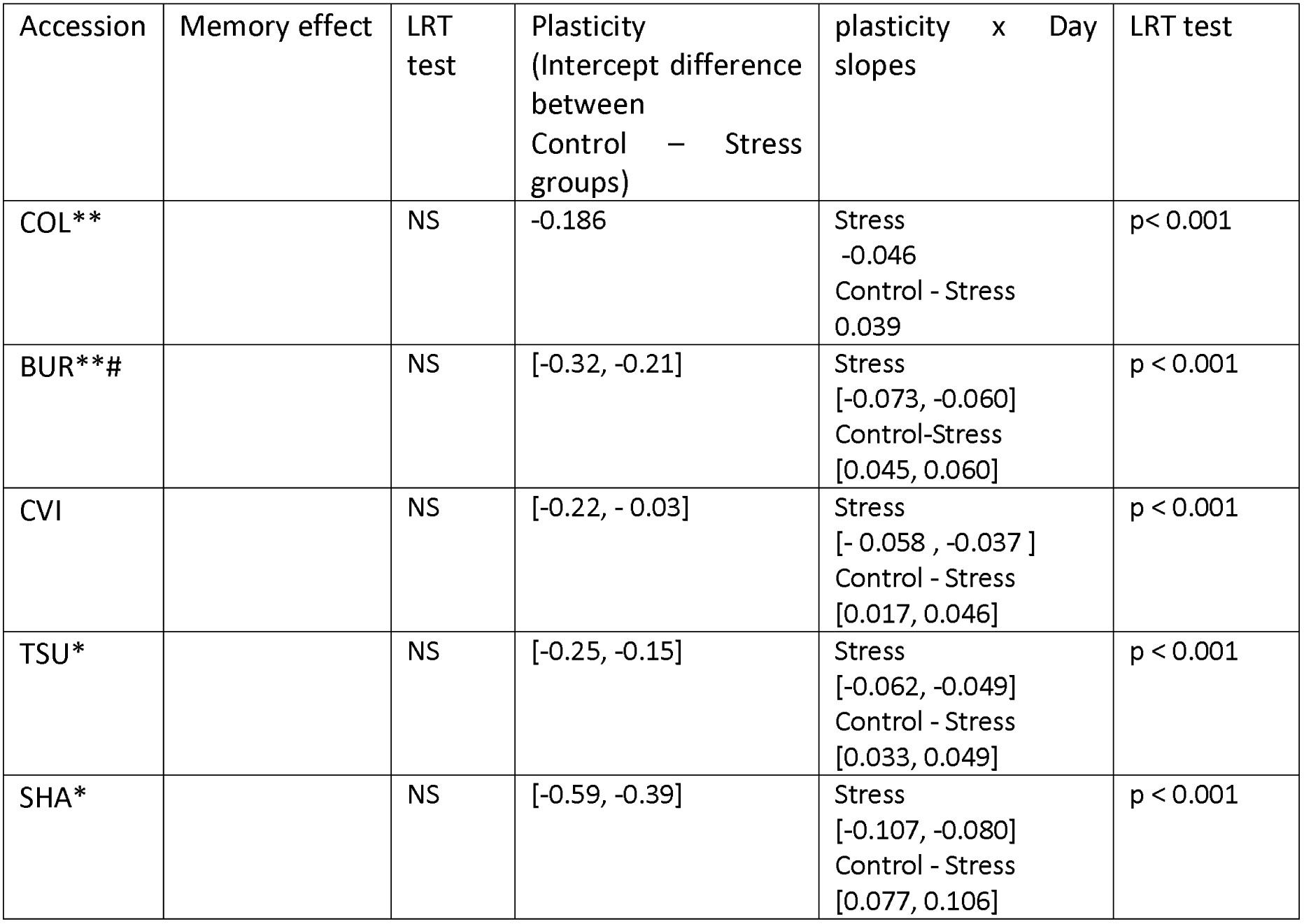
Effects of environmental states in generations G1/G2 and P4 on the mean relative growth rate (RGR) from days 13 to 16 in P4. Results from linear mixed model analysis and model after selection. In the table, accessions with random effects that differ in line/individual variance between the two G1/G2 treatment levels are indicated by symbol "**". Symbol "*" indicates accessions where the variance between lines is retained in the model. "#" indicates accessions for which the pot effects (linear effect of pot number on the robot) were retained.

**Table 3.**
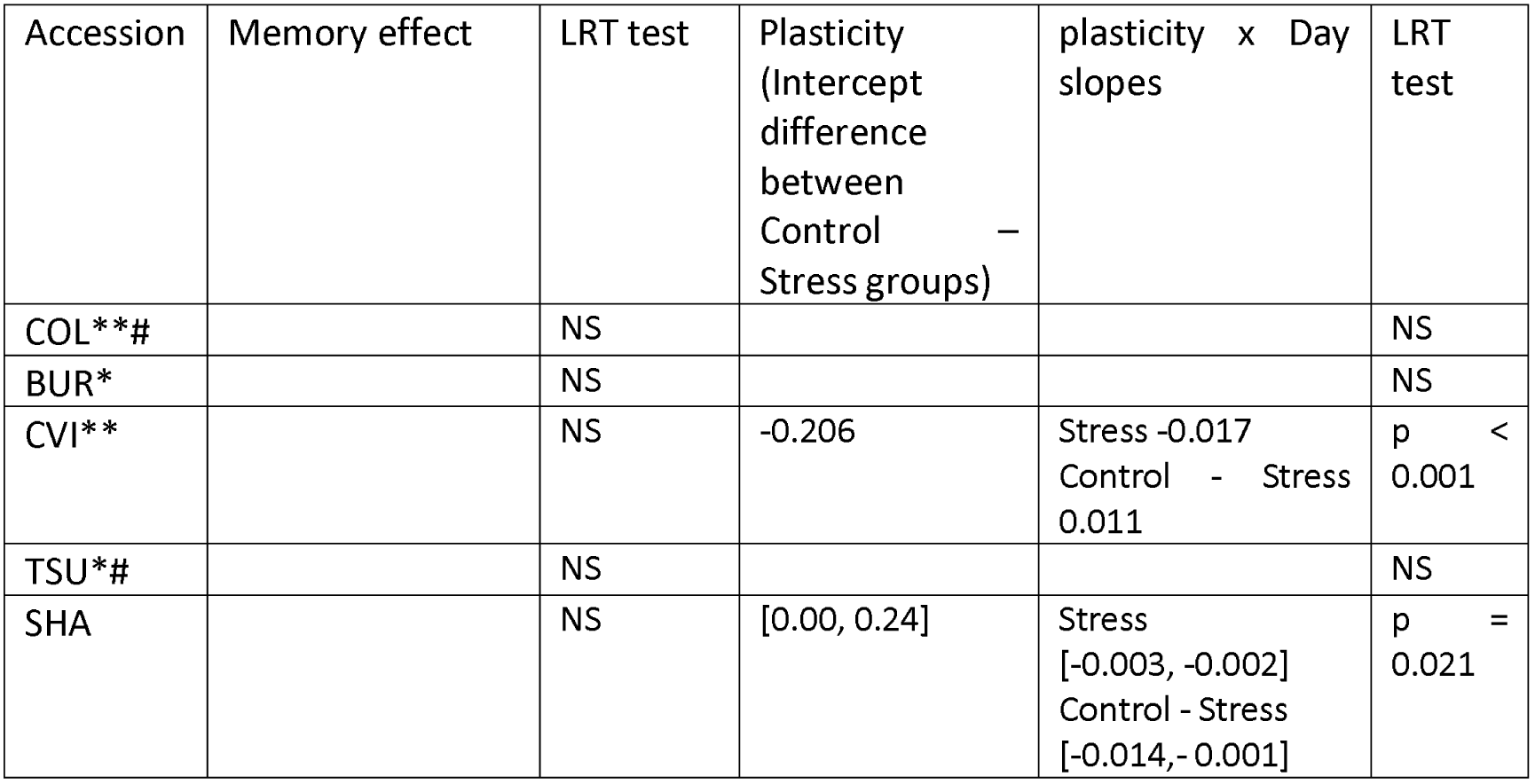
Effects of environmental states in G1/G2 and P4 on the mean relative growth rate from days 25 to 28 in P4. Results from linear mixed model analysis. In the table, accessions with random effects that differ in line/individual variance between G1/G2 groups are indicated by "**". Symbol "*" indicates accessions where the variance between lines is retained in the model. "#" indicates accessions for which the pot effects (linear effect of pot number on the robot) were retained.

### Limited presence and persistence of maternal trait-based effects

Models with maternal trait effects on individuals in P4 find in two out of 125 tests that the slope of the maternal trait regression depends on the environmental regime experienced by ancestors in G1/G2 (Supporting Information Table S2) and in two cases on the environmental regime in P4. By inspecting the models per accession, we can make an assessment of whether maternal trait slopes changed by mild stress would affect the persistence of these maternal effects. This can be the case when slopes of traits on themselves have become larger in absolute value or when a causal chain of traits has obtained a stronger weight (Kirkpatrick & Lande, 1989). Only for compactness and in Col and Bur do we find a maternal trait-based effect where the trait has an effect on itself. While for Col-0 the slope of the maternal effect is changed in plants from lines that have experienced stress in previous generations, this is not the case for Bur (Supporting Information Table S2). There are no enchained maternal trait effects that would lead to lagged responses over several generations.

When we inspect the maternal trait dependency (Fig. 4) of compactness in Col, we note that the ranges of maternal trait values differ between descendants of stress and control treatments in G1/G2. The results do show that maternal trait models have slopes that can depend on historical and current environments and that the scope for a change in the persistence of maternal effects due to mild stress is limited (Fig. 4). Indeed, the slopes of the effects are not particularly strong, nor general across accessions and they do not seem a valid candidate for persistent transmission of trait variation. Only 5 out of 125 tests for effects of maternal traits on offspring were significant and had interpretable confidence intervals, which is very close to the type I error rate. We therefore conclude that the number of heritable effects transmitted through maternal trait-based effects is negligible.

**Fig. 4.**
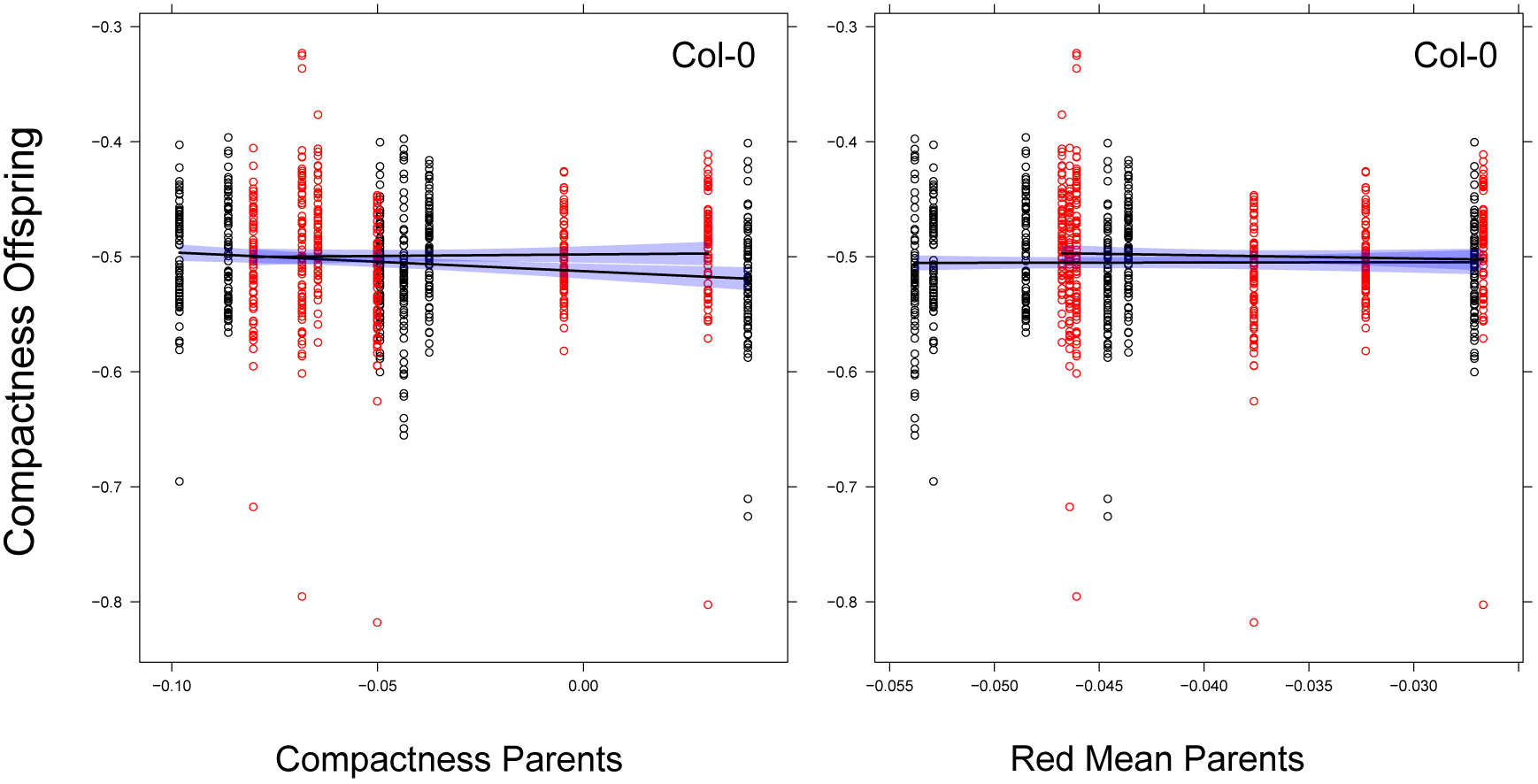
Effects of maternal traits in Col-0. Dependencies of offspring trait values on maternal trait values are shown for log-transformed rosette compactness in dependence on two maternal trait values. In the mixed model analysis, we detected significant effects of maternal compactness but not of maternal Mean Red value. Linear regressions are estimated per ancestral environment. Data points are red for individuals with ancestors under stress in G1/G2, black for individuals with ancestors under control.

In 13 out of 25 trait × accession models, individual variation is enlarged in the stress environment in P4 (Supporting Information Table S3), in 5 out of 25 models the variation between individuals is larger among descendants of individuals stressed in G1/G2. In two cases this variance is smaller.

### Mild drought induces DNA methylation variation and uncorrelated transcriptome changes

To complement the phenotypic analysis, we investigated the impact of mild drought on genomic DNA methylation patterns using the reference accession Col-0, which shows the strongest phenotypic response (Fig. 2b) and for which a wealth of epigenomic data are available. WGBSeq was performed on DNA extracted at day 29 from leaves of pools of treated and control plants at P4 (C1C2C3 descendants, Fig 1; Supporting Information Tables S4 and S5). Overall, cytosine methylation levels are similar between control- and stress-treated leaves (Supporting Information Fig. S1a), albeit slightly higher than in previous reports (8.6% and 9% of methylated cytosines vs. 6.7%), presumably because of differences in mapping and methylation calling methods as well as in the organs examined (Cokus *et al.*, 2008; Lister *et al.*, 2008). Other global measures, such as the distribution of methylation between the three types of sites and annotations are also identical for control- and stress-treated leaves (Supporting Information Fig. S1b and c). Thus, we conclude that mild drought does not directly affect overall DNA methylation patterns in Arabidopsis.

To identify local differences, methylation levels were compared at individual cytosine positions as well as in 100 bp windows for each of the three types of sites (CG, CHG and CHH) separately (see Methods). Based on this approach, we could identify 286 differentially methylated positions (DMPs) and 1360 differentially methylated regions (DMRs), most of which are defined by single 100 bp windows (Supporting Information Tables S6 and S7). All DMPs map to CG sites whereas most DMRs (95%) are CHH DMRs only (Fig. 5a). The vast majority of CG DMPs (93%) are within methylated gene bodies (Fig. 5a and b) and they reflect almost equally either increased or decreased methylation levels in treated plants compared to controls, consistent with the notion that gene body methylation tends to vary stochastically across generations at individual CG sites (Becker *et al.*, 2011; Schmitz *et al.*, 2011; Jiang *et al.*, 2014). On the other hand, CHH-DMRs are mainly located over TEs sequences and tend to reflect hypermethylation in treated plants (Fig. 5c and d, Supporting Information Fig. S2).

**Fig. 5.**
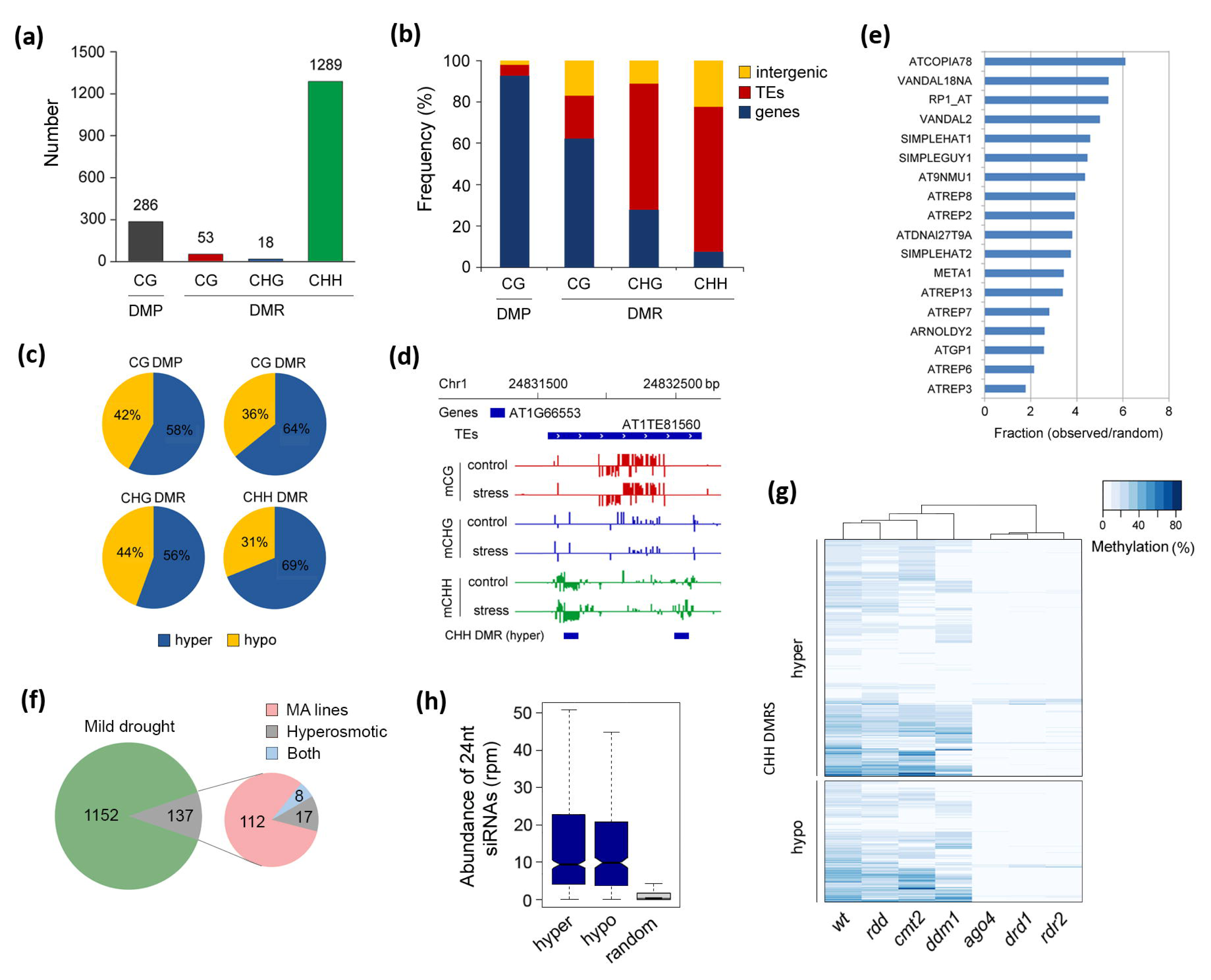
Characterization of stress-induced local DNA methylation changes. (a) Number of DMPs and DMRs for each sequence context (CG, CHG and CHH). (b) Annotation of DMPs and DMRs in relation to genes, TEs and intergenic regions. (c) Distribution of local gains and losses of DNA methylation across DMPs and DMRs. (d) Example of CHH DMRs on a TE. (e) Graphical representation of the 18 TE families that show more DMRs than expected by random (*p*-value < 0.01). (f) Overlap (including 500bp flanking windows) of DMRs induced by mild drought and DMRs found in mutation accumulation (MA) lines (Becker *et al.*, 2011; Schmitz *et al.*, 2011) or induced by hyperosmotic stress (Wibowo *et al.*, 2016) (g) Hierarchical clustering based on average CHH methylation levels in wild-type (wt) and mutants for the RdDM (*rdr2*, *ago4* and *drd1*), CMT2 (*ddm1* and *cmt2*) and DNA demethylation (*rdd*) pathways in regions overlapping hyper or hypermethylated CHH-DMRs. (h) Abundance of 24nt siRNAs in random or CHH-DMR containing TEs.

As different TE families may show different sensitivity to environmental cues (Pecinka *et al.*, 2010; Yu *et al.*, 2013; Grandbastien, 2015; Matsunaga *et al.*, 2015; Quadrana *et al.*, 2016), we assessed whether CHH-DMRs are preferentially localized over specific TE families. Out of the 326 TE and other repeat families annotated in the TAIR10 Arabidopsis genome, 164 show at least one DMR and 18 families are enriched in DMRs compared to the random expectation (Fig. 5e). These include the LTR-retrotransposon family *ATCOPIA78*, which is known to be sensitive to biotic and abiotic stress (Yu *et al.*, 2013; Quadrana *et al.*, 2016; Matsunaga *et al.*, 2015). On the other hand, only a small percentage of CHH DMRs caused by mild drought overlap with DMRs that arose spontaneously in mutation accumulation lines (9.3%;Hagmann *et al.*, 2015) or that were induced by hyperosmotic stress (1.9%;Wibowo *et al.*, 2016) (Fig. 5f). Thus, we conclude that mild drought induces a limited number of robust DNA methylation changes over regions that are distinct from those subjected to stochastic or salt-induced DNA methylation variation.

To investigate further the CHH-DMRs induced by mild drought, we compared their CHH methylation level in different DNA methylation mutants (Stroud *et al.*, 2013) and found that most correspond to regions targeted by the RNA-directed DNA methylation (RdDM) pathway, which involves the DNA methyltransferase DRM2, rather than by the alternative CHH maintenance methylation pathway mediated by the DNA methyltransferase CMT2 (Fig. 5g). Consistent with these findings, TE sequences overlapping drought induced CHH-DMRs have a high abundance of matching 24nt small RNAs (Fig. 5h). Moreover, no correlation was detected between drought induced CHH-DMRs and regions subjected to active DNA demethylation (*rdd* mutant; Fig.5g). In conclusion, mild drought directly affects mainly sequences targeted by RdDM.

To determine if genes with changes in DNA methylation near or within them are drought stress responsive, we performed RNA-seq on leaves isolated from Col-0 plants directly exposed to mild drought or control treatments and grown for 3 weeks on the Phenoscope. Consistent with the results of previous studies of the transcriptional response to mild drought stress (Cubillos *et al.*, 2014; Clauw *et al.*, 2015), differential analysis of the two RNA-seq datasets identified significant changes in steady state mRNA levels for 468 genes (FDR < 0.05, >0.5 log2 fold change, 205 and 263 genes with lower and higher expression under mild drought, respectively; Supporting Information Table S8), but not for any of the annotated TE sequences. Gene Ontology analysis revealed enrichment for several stress-related categories, including response to water deprivation, for which the highest significance level was observed. However, none of the genes known to be involved in DNA (de)methylation appeared to be affected by mild drought, which leaves the question open as to which factors induce DNA methylation changes during mild drought.

Among the 468 genes detected as transcriptionally responsive to mild drought in our conditions, only two are affected by CG DMPs, which are likely inconsequential given the lack of function associated with gene body methylation. Another two are located less than 500bp from a DMR (Supporting Information Fig. S3a). These two DMRs are of the CHH type but do not correspond to annotated TE sequences. One DMR maps to the promoter region of gene *AT5G35735,* which encodes an auxin responsive protein of unknown function. The other DMR is located within the first intron of gene *AT3G10340*, which encodes a putative phenyalanine ammonia-lyase that may be involved in plant defense against biotic and abiotic stresses (Raes *et al.*, 2003). Given the first intron large size (1.3kb), it likely contains regulatory sequences (Morello & Breviario, 2008). Moreover, hypermethylation of the promoter DMR of *AT5G35735* and hypomethylation of the intronic DMR of *AT3G10340* in response to mild drought are associated with down and up regulation, respectively (Supporting Information Fig. S3b). Taken together, these findings suggest a causal link between altered DNA methylation and altered gene expression for these two genes. However, the observation that most genes affected by mild drought are not proximal to drought-induced DMPs or DMRs indicates that changes in DNA methylation have a marginal role in the phenotypic response of plants to mild drought.

### DNA methylation variation is not transmitted to the progeny

Finally, we tested whether the DNA methylation changes directly induced by mild drought could be transmitted from stressed plants to their progeny. Taking into consideration the possibility of cumulative effects over successive generations of growth under stress, G3 progenies of Col-0 C_1_C_2_ and S_1_S_2_ plants were chosen for further analysis (Fig. 1a). WGBseq was performed on DNA extracted from unstressed individuals of the progeny of five Col C_1_C_2_ and S_1_S_2_ founder lines (Methods and Supporting Information Tables S4 and S5). Differential DNA methylation was investigated as described above using the five C_1_C_2_ or S_1_S_2_ progenies as biological replicates. Following this approach, no single consistent DMR could be identified between the two types of progeny. However, there was a marginal increase in the amount of stochastic variation in DNA methylation for the three sequence contexts among progenies derived from the five stressed parental lines (Supporting Information Fig. S4). In conclusion, our results suggest that targeted and specific DNA methylation changes induced by mild drought are not transmitted to the next generation, although stress exposure may increase methylome heterogeneity among progeny of stressed plants.

## Discussion

It has been proposed that exposure to environmental cues can trigger phenotypic changes that are inherited for more than one generation, and that this occurs through epigenetic mechanisms (Bossdorf *et al.*, 2008; Richards *et al.*, 2017). In this study, we showed that water deficit applied before the reproductive stage in two successive generations affects the vegetative growth of individuals negatively. Memory effects of mild drought stress after two successive generations of exposure are limited to changes in amounts of individual variation in phenotypic traits, but the variance can be increased or decreased depending on the accession and trait considered. Furthermore, although changes in DNA methylation in response to mild drought were observed, these were only marginally associated with changes in gene expression and were not transmitted to the immediate progeny of the affected individuals, even after two successive generations of exposure to stress. In other words, there is no substantial evidence that drought stress would modify the heritability of methylation patterns and of maternal trait-based effects. Therefore, our results add to the growing body of evidence against transgenerational epigenetic variation being a common response of plants to changes in the environment.

### Intergenerational plasticity is limited and does not lead to transgenerational effects

Stresses consistently affect the expression of a large number of genes and induce in a number of cases CHH hyper or hypomethylation of a variable number of TE sequences and other repeat sequences (for example Dowen *et al.*, 2012; Eichten & Springer, 2015; Secco *et al.*, 2015; Wibowo *et al.*, 2016; this study). However, gene expression changes rarely correlate with DNA methylation changes (Meng *et al.*, 2016) and the extent of the latter as well as the mechanisms at play may differ radically between plant species for a given stress (Secco *et al.*, 2015). Furthermore, DNA methylation changes may or may not be transmitted to the next generation, for reasons that are unclear. Thus, while in Arabidopsis many of the CHH-DMRs induced by salt stress are transmitted to the immediate progeny (Wibowo *et al.*, 2016), this is not the case for the CHH-DMRs induced by mild drought (this study) and in rice there was also no transmission of the CHH-DMRs induced by phosphate starvation (Secco *et al.*, 2015). Thus, evidence so far points to a clear effect of environmental factors in triggering DNA methylation changes, which however do not persist across more than one generation. Indeed, different mechanisms preventing transgenerational transmission of environmentally induced epigenetic states have been described (Baubec *et al.*, 2014; Crevillen *et al.*, 2014; Iwasaki & Paszkowski, 2014). Nonetheless, true transgenerational epigenetic variation exists in nature (Quadrana & Colot, 2016) and what generates it remains unresolved. Analyses of natural populations are now just beginning to investigate this question, with no clear answer so far, except that most DNA methylation variants seen in nature are likely caused by DNA sequence variation and therefore are by definition not truly epigenetic (Schmitz *et al.*, 2013; Li *et al.*, 2014; Dubin *et al.*, 2015; Kawakatsu *et al.*, 2016b; Niederhuth *et al.*, 2016; Quadrana *et al.*, 2016).

At the morphological level, we detected different responses between accessions, with both plasticity and some trait-based maternal effects playing a role. Maternal trait-based effects with potentially lasting effects occur but for rosette compactness only. Such effects should dampen quickly when selection does not favor increased compactness for example and these maternal effects are potentially swamped by environmental variability. Drought stress changes trait variances but not slopes of maternal trait-based effects. No new significant slopes were created. Therefore mild drought stress did not change this presumed mechanism of non-genetic heritability.

### Plasticity is likely adaptive, a memory effect not

Transgenerational effects are often presumed to be adaptive. Models show that environmentally induced epiallelic variation can be favored over purely stochastic switching (Furrow & Feldman, 2014). The general lack of such effects in our experiment might indicate that they are actually unnecessary in the range of imposed environments. The remaining responses observed would then rather be of a non-adaptive nature and reflect mechanistic constraints in plant development. In agreement with models of adaptation (Kuijper & Hoyle, 2015), we find that responses by means of phenotypic plasticity are stronger than by maternal effects. The changes in trait variance could indicate that mild drought stress rather affects the predictability of the near future, without much of a consistent trend in expectations. Our observed changes might then be more in support of a bet-hedging strategy (e.g., Crean & Marshall, 2009).

Modeling suggests a potential for DNA methylation-based transgenerational epigenetics to endow plants with a means to generate adaptive heritable phenotypic variation in response to changing environments (Bossdorf *et al.*, 2008; Geoghegan & Spencer, 2013b; Geoghegan & Spencer, 2013a; Uller *et al.*, 2015; Kronholm & Collins, 2016). We did not detect this. The absence of a clear change in maternal-trait based effects seems to indicate that mild drought stress does not induce strong physiological changes that would affect transmission of information or resources. Similarly, we observed that relative growth rates respond to this stress in a transient manner, clearly affecting final sizes but with return to control-like relative growth rates after stress treatment. It has been proposed to see phenotypes as weighted sums of ancestral contributions (Sultan, 2017). Similarly to Tallis’ vision of ancestral genotypic regression (Tallis, 1987), phenotypic plasticity would be broadened to include effects of ancestral environments. Maternal trait-based effects are such ancestral genotypic × environmental regressions. We find that contributions of ancestral states were limited and decay quickly.

In conclusion, our study provides strong support to the notion that plants first respond to physiological stresses through well-defined and conserved transcriptional networks (Juenger, 2013) (Ding *et al.*, 2014; Clauw *et al.*, 2015) or immediate parental influences on offspring phenotypes (Herman & Sultan, 2011; Wibowo *et al.*, 2016). Whether transgenerational epigenetic variation in nature is caused by more dramatic environmental conditions than those tested so far in the laboratory, or by combinations of several mild stresses, or by mutations in genes, such as Arabidopsis *DDM1*, that are involved in the epigenetic control of TE remains to be determined (Quadrana & Colot, 2016).

## Acknowledgments

We thank members of the Colot lab for discussions. This work was supported by funding from the Agence Nationale de la Recherche (project MEMOSTRESS, grant n^0^ ANR-12-ADAP-0020-01 (to V.C, O.L. and T.J.M.V.D.) and the European Commission Framework Programme 7, ERC Starting Grant ‘DECODE’ / ERC-2009-StG-243359 (to O.L). The IJPB benefits from the support of the LabEx Saclay Plant Sciences-SPS (ANR-10-LABX-0040-SPS). ABS was supported by a postdoctoral fellowship from the Brazilian National Council for Scientific and Technological Development (CNPq – Brazil). LQ was recipient of postdoctoral fellowships from the ANR-10-LABX-54 MEMOLIFE and ANR-11-IDEX-0001-02 PSL Research University.

## Author contributions

T.J.M.V.D., A.B.S., O.L., and V.C. planned and designed research; A.B.S., E.G., A.M., L.B and S.T. performed experiments and collected the data; T.J.M.V.D., A.B.S., L.Q., analyzed the data; J.J.G. contributed unpublished results; T.J.M.V.D., A.B.S., O.L. and V.C. wrote the manuscript with the help of all authors.

## Supporting Information legends

**Fig. S1**. Genome-wide DNA methylation patterns are similar between leaves of stressed and non-stressed plants.

**Fig. S2**. Chromosomal distribution of local gains and losses of DNA methylation found between leaves of stressed and non-stressed plants.

**Fig. S3**. Differential methylation is weakly correlated to changes in gene expression in response to mild drought.

**Fig. S4**. The progenies of stressed lines exhibit increased methylome instability.

**Table S1**. Effects of environmental state in G1 and G4 on the residual variance in initial log(PRA), projected rosette area.

**Table S2**. Estimates of trait-based maternal effects per accession.

**Table S3.** Estimates of Individual within-line variances in models with trait-based maternal effects per accession.

**Table S4.** Summary statistics of whole genome bisulfite sequencing data.

**Table S5.** Total fraction of methycytosines and distribution in each sequence context

**Table S6.** List of differentially methylated positions.

**Table S7.** List of differentially methylated regions.

**Table S8.** List of differentially expressed genes.

